# IL-17^+^ mast cell/T helper cell axis in the early stages of acne

**DOI:** 10.1101/2021.06.29.450328

**Authors:** Yoan Eliasse, Edouard Leveque, Lucile Garidou, Louise Battut, Brienne McKenzie, Thérèse Nocera, Daniel Redoules, Eric Espinosa

**Author notes:** Correspondence: Eric Espinosa, INSERM U1037, CRCT, 2 Avenue Hubert Curien, CS 53717, 31037 Toulouse cedex 1 - France.

## Abstract

Acne is a multifactorial disease driven by physiological changes occurring during puberty in the pilosebaceous unit (PSU) that leads to sebum overproduction and a dysbiosis involving notably *Cutibacterium acnes*. These changes in the PSU microenvironment lead to a shift from a homeostatic to an inflammatory state. Indeed, immunohistochemical analyses have revealed that inflammation and lymphocyte infiltration can be detected even in the infraclinical acneic stages, highlighting the importance of the early stages of the disease. In this study, we utilized a robust multi-pronged approach that included flow cytometry, confocal microscopy, and bioinformatics to comprehensively characterize the evolution of the infiltrating and resident immune cell populations in acneic lesions, beginning in the early stages of their development. Using a discovery cohort of 15 patients, we demonstrated that the composition of immune cell infiltrate is highly dynamic in nature, with the relative abundance of different cell types changing significantly as a function of clinical lesion stage. Within the stages examined, we identified a large population of CD69^+^ CD4^+^ T cells, several populations of activated antigen presenting cells, and activated mast cells producing IL-17. IL-17^+^ mast cells were preferentially located in CD4^+^ T cell rich areas and we showed that activated CD4^+^ T cells license mast cells to produce IL-17. Our study reveals that mast cells are the main IL-17 producers in the early stage of acne, underlying the importance of targeting the IL-17^+^ mast cell/T helper cell axis in therapeutic approaches.

## INTRODUCTION

Acne vulgaris is a chronic inflammatory skin disease that appears concomitantly with hormonal changes at puberty and constitutes the most common cutaneous disorder in adolescents and young adults. It affects the skin pilosebaceous units, which consist of the hair shaft and the hair follicle with an attached sebaceous gland. Excess sebum and hyperkeratinisation plug up the pore, leading to dramatic changes in the PSU microenvironment (Tuchayi et al., 2015).

The pathogenesis of acne is not fully understood. Hormonal stimuli at puberty induce sebocyte and ductal keratinocyte activation and proliferation in the PSU, leading to hyperproduction of sebum and disturbed keratinization (Tuchayi et al., 2015). Sebum accumulation and progressive clogging of the pore create a new environment that impacts the PSU microbiota and notably foster *C. acnes* proliferation. Ensuing cellular stress and disruption in homeostasis provide several pro-inflammatory cues. These changes, in combination with other factors that remain to be elucidated, drive a naive PSU into the acne cycle (Saurat, 2015) starting with the microcomedone stage. Microcomedones are histological observations, invisible by eye, that can be detected by skin surface biopsy (Cunliffe et al., 2004). Next the microcomedone evolves into a closed comedone (CC or whitehead) or open comedone. These stages are associated with a lack of visible inflammation. While some lesions remain as comedones and resolve, others evolve toward an exacerbation of the inflammatory process leading to the first visible inflamed lesions named papules (PA). These latter lesions can regress or evolve toward pustule and subsequently nodule and/or cyst (Do et al., 2008, Saurat, 2015, Tuchayi et al., 2015)

The early changes that occur inside a PSU and underlie the progression through the CC stage and next through the PA stage are poorly understood. Notably, the role of the skin immune system (SIS) remains subject to much debate. The early immune response observed in acne is only partially described and appears to be different from the classical inflammatory response mounted against a pathogen, in which the early steps are dominated by neutrophil and subsequently monocyte infiltration upon danger/pathogen recognition. Pioneering histology studies of skin biopsies reported that T lymphocytes were the first cells to infiltrate early lesions (Jeremy et al., 2003, Layton et al., 1998, Norris and Cunliffe, 1988). Moreover, some T cells infiltrating early stages of acne were found to be specific for *C. acnes* antigens (Mouser et al., 2003). A consensus has now developed suggesting that the inflammatory response observed after the PA stage first emerges in the very early stages of the disease (Do et al., 2008) and that inflammation is the starting point of the acne process. Nevertheless, the immune surveillance system of the skin is complex (Belkaid and Tamoutounour, 2016, Kabashima et al., 2019) and several resident sentinel cells (dendritic cells, macrophages and mast cells) might be involved in perceiving the alarm signals and in starting the early response. Mast cell are particularly abundant in skin and mucosa and are strategically located near blood vessels and nerve endings (Valent et al., 2020). They can produce a large array of mediators and respond to perturbations in their environment, leading to their involvement in several inflammatory disorders (Siiskonen and Harvima, 2019, Valent et al., 2020). Whilst the role of mast cells in acne is poorly understood, these cells are known to play important roles in the skin immune system (Voss et al., 2021). Furthermore, different resident and recirculating T cell subsets are also known to participate in immune surveillance of the skin (Watanabe et al., 2015) and might be involved in the initiation of the immune response in acne. *C. acnes* is an extracellular pathogen expected to induce a type 3 immune response orchestrated mainly by Th17 cells (Annunziato et al., 2015). IL-17 and related cytokines are proinflammatory cytokines known to promote anti-microbial protective responses and barrier maintenance but also pathogenic inflammation upon chronic activation (McGeachy et al., 2019). IL-17 (Ebrahim et al., 2019) and Th17 cells (Agak et al., 2014, Kelhala et al., 2014) were reported to be associated with acne pathogenesis but their functional role in acne remains understudied.

To gain a more comprehensive understanding of the early inflammatory response in lesions, we profiled the evolution of skin immune cell populations during the early stages of acne (CC and PA). Using a comprehensive global approach combining flow cytometry, confocal microscopy, and bioinformatics, we highlighted that multiple immune cell populations are swiftly recruited and that the evolving nature of these immune populations is a defining feature of lesion stage. Microscopy approaches allowed us to identify that mast cells were the main IL-17 producers in the CC stage and that mast cell IL-17 production was licensed by activated CD4^+^ T cells. Our study thus provides unprecedented insight into the immune events that shape lesion development and identify potential targets for therapeutic intervention.

## MATERIALS AND METHODS

### Skin biopsies

15 young male adults with acne on the back (phototype II to IV), presenting Investigator Global Assessment score ≥ 3, were enrolled in the study. Main exclusion criteria included: presence of any skin condition that would interfere with the diagnosis or assessment of acne vulgaris, sun exposure, anti-inflammatory treatments, topical acne or antibiotic treatments within 14 days prior to baseline, use within 1 month prior to baseline of systemic antibiotics, use within 6 months prior to baseline of oral retinoids. Patients gave written informed consent under a protocol approved by local ethic committee (CCP Sud-ouest et Outremer III, #Id RCB: 2015-A01858-41) an in agreement with the Declaration of Helsinki Principles. Punch biopsies (3 biopsies of 4mm and 2 biopsies of 3 mm diameter) were taken from the back of the patients (Supplementary Figure S1). Biopsies were stored in buffer (MACS Tissue Storage Solution, Miltenyi Biotec) at 4°C and processed within 2 hours for cell isolation or PFA fixation.

### Flow cytometry

Cells from whole skin were dissociated into single-cell suspension by combining mechanical dissociation and enzymatic digestion. Briefly, the biopsies were washed in two successive PBS bathes, cut into 4 pieces and enzymatically digested by using the whole skin dissociation kit from Miltenyi biotec without enzyme P for 5 hours at 37°C under agitation (500 rpm). 500μL of cold RPMI 10% SCF were added to the samples and mechanical dissociation was achieved by using the BD™ Medimachine System (Becton-Dickinson). We optimized the tissue disaggregation in our experimental settings as follows: the 4 pieces were next placed into a disposable disaggregator Medicon™ with 50 μm separator mesh in 1 mL of ice-cold RPMI 10% SCF and processed in the Medimachine System by using a disaggregation time of 40 sec. The cell suspension was recovered from the Medicon unit with a 5-mL disposable syringe (the chamber was rinsed twice to increase yield) and was filtered (70 μm Filcon™ disposable filter device) and washed twice with RPMI 10% FCS. Cells were next split into three Eppendorf tubes and used for flow cytometry

Antibodies were mixed according to the 2 multicolor flow cytometry panels (P1 and P2) at the concentration advised by the manufacturers in PBS 10% human serum (Supplementary Figure S1 and Table S1) plus viability dye (Fixable viability dye-eFluor 506, eBiosciences). Cells were stained for 30 minutes at 4°C (with P1, P2 Ab mixes or PBS) washed twice in PBS/EDTA and acquired using BD LSR fortessa, flow rate calibration method was used to calculate absolute cell numbers. Data were analyzed using FlowJo software (V10, Tree Star).

### Biopsies - Immunofluorescence – confocal microscopy

Biopsies for microscopy analysis were washed twice in PBS and fixed in PBS 4% PFA overnight at 4°C. The biopsies were washed for 5 min in a first bath of PBS and then for 24h in a second bath of PBS at 4°C and next bathed in PBS 30% sucrose for 48h at 4°C. Biopsies were next rinsed in two successive bathes of PBS for 2 hours and embedded in OCT™-Compound (Optimal Cuting Temperature-Compound; Tissu-Tek® Sakura), frozen and stored at −80°C until use. Biopsies were cut into 30 or 40 μm sections with a cryostat (Microm HM550 from Thermo Scientific) and mounted on adhesive glass slides and stored at −80°C until further use. Frozen tissue sections were incubated for 10 minutes in PBS at 20°C and processed for antigen retrieval in HIER-EDTA pH 9.0 buffer (Zytomed Systems) at 90°C for 15 minutes. They were rinsed in PBS and blocked with PBS 20% human serum 0.3% saponin and next incubated with primary Abs diluted in PBS 20% human serum 0.3% saponin (Supplementary Table S1) for 16 hours at 4°C in a humidified chamber. They were next washed three times for 10 min and incubated with secondary Alexa Fluor-conjugated Abs (Invitrogen) for 2 hours at room temperature. After 3 washes of 10 minutes, nuclei were stained with DAPI for 5 minutes at 20°C. Images were acquired using Zeiss LSM 710 or LSM 780 confocal microscope (Zeiss, Oberkochen, Germany) and Zen Black software with a 40X/1.4 Oil Plan-Apochromatic objective.

### Principal component analysis (PCA) and data clustering

We performed hierarchical clustering on flow cytometry data (numbers of the different cell population identified and percentages of Th cell subsets measured for the 3 stages of acne). Data were scaled and clustered, heatmap was obtained using heatmap.2 function of the gplots R package (complete linkage clustering using a Euclidean distance measure). PCA was next applied on these flow cytometry data to reduce the dimensionality (using the FactoMineR R package). The individuals’ graph showed the distance between observations in the PC1 vs PC2 scatter plot and the variables’ graph (correlation circle) showed the correlation among variables.

### Primary human mast cell lines

Peripheral blood mononuclear cells (PBMCs) were obtained from buffy coats (Etablissement Français du Sang). CD34+ precursors cells were isolated from PBMCs (EasySep™ Human CD34 Positive Selection Kit, STEMCELL Technologies). CD34+ cells were grown under serum-free conditions using StemSpan™ medium (STEMCELL Technologies) supplemented with recombinant human IL-6 (50 ng mL-1; Peprotech), human IL-3 (10 ng mL-1; Peprotech) and 3% supernatant of CHO transfectants secreting murine SCF (a gift from Dr. P. Dubreuil, Marseille, France, 3% correspond to ~50 ng.mL^−1^ SCF) for one week. Cells were grown in IMDM Glutamax I, sodium pyruvate, 2-mercaptoethanol, 0.5% BSA, Insulin-transferrin selenium (all from Invitrogen), ciprofloxacin (10 μg.mL^−1^; Sigma Aldrich), IL-6 (50 ng.mL^−1^) and 3% supernatant of CHO transfectants secreting murine SCF for 8 weeks and tested both phenotypically (Tryptase^+^, CD117^+^, FcεRI^+^) and functionally (β-hexosaminidase release in response to FcεRI crosslinking) before use in experiments. Only primary cell lines showing more than 95% CD117^+^/FcεRI^+^ cells were used for experiments.

### Mast cell-CD4^+^ T cell cocultures - Immunofluorescence – confocal microscopy

After 48h coculture, cells were transferred on poly-L-lysine-coated slides and then fixed with 4% paraformaldehyde in PBS. Cells were first permeabilized and blocked in 10% normal human serum in PBS containing 0.3% saponin. Cells were stained with the following primary antibodies in PBS containing 1% BSA, 0.3% saponin for 2 hours at room temperature: anti-CD4, anti-IL-17A and anti-tryptase Abs (Supplementary Table S1). After washing, matched secondary antibodies (Alexa Fluor-conjugated donkey Abs, Invitrogen) were applied in PBS containing 1% BSA, 0.3% saponin for 45 min at RT. After 3 washes of 5 minutes, nuclei were stained with DAPI for 5 minutes at 20°C. The samples were mounted and examined using a Zeiss LSM 780 confocal microscope with a 63× Plan-Apochromat objective (1.4 oil). Scoring of the slides was performed in a blinded fashion by evaluating for each condition at least 200 mast cells in randomly selected fields from 3 independent experiments.

### Real-time quantitative PCR

Total RNA was extracted from hMCs by using the phenol-chloroform method. IL-17A gene expression levels were measured by quantitative RT-PCR with Power SYBR Green technology (Applied Biosystems). Probes were obtained from Applied Biosystems as Assays-on-Demand™ Gene Expression Assays (glyceraldehyde-3-phosphate dehydrogenase GAPDH: Hs02758991_g1, IL-17A: HS 00174383_m1. Changes in relative gene expression were calculated using the 2^−ΔΔCT^ method with normalization to GAPDH mRNA. All reactions were performed in three independent assays with three technical replicates.

### Mast cell-CD4^+^ T cell cocultures

2.10^5^ CD4^+^ effector/memory T cells (isolated from healthy donor PBMCs by negative selection using EasySep™ Human Memory CD4 T Cell Enrichment Kit, Stemcell) were incubated with 2.10^5^ primary human mast cells (cell line established from a different donor) with or without anti-CD3/CD28-coated beads (Dynabeads™, Gibco) at a T cell:bead ratio of 10:1 in RPMI 10% FCS supplemented with 20 ng.mL^−1^ SCF. For RT-qPCR experiments, cells were harvested after 4, 24 or 48h incubation, stained with anti-CD117-PECy7 mAb (5 μg.mL^−1^) and viability dye and sorted by a FACSMelody™ cell sorter (BD biosciences). For immunofluorescence experiments, cells were transferred on poly-L-lysine-coated slides after 48h coculture.

### Statistics

GraphPad Prism software (V.9; GraphPad Software, La Jolla, CA, USA) and R 4.0.3 were used for statistical analysis (stats package). Differences between groups were analyzed using one-way ANOVA (Dunnett’s Multiple Comparison Test) or Friedman test with Dunn’s post hoc test. Principal component Analysis (PCA) was applied on flow cytometry data (variables gathered in the heatmap figure 3a) with the FactoMineR R package.

## RESULTS

### Immune cell landscape of the early stages of acne

To quantify the immune infiltrate of the first stage of acneic lesion development and provide a detailed cartography of the spatial relationship between immune cell populations, we used both flow cytometry and microscopy approaches to study skin biopsies from a cohort of 15 young adults presenting with mild to moderate acne on the back (Figure S1a-b). We focused our study on the early stages of pathology by collecting 3 types of skin biopsies: uninvolved (UI, i.e. skin without lesion in acne patient) skin, closed comedone (CC) and papule (PA) for each patient (Figure S1a,c). We developed a protocol for cell dissociation from skin biopsies that maximizes viable immune cell recovery (notably macrophages and mast cells that tend to be attached to the extracellular matrix) and minimizes cell surface marker digestion during tissue preparation. We devised two surface maker panels in order to analyze the main leukocyte populations by multicolor flow cytometry (Figure S1). The first panel was designed to identify lymphocytes and mast cells (Figure S2). The second panel was designed to identify the monocyte/dendritic cell lineage populations and neutrophils (Figure S3). Frequencies and composition of the cell populations identified by flow cytometry are presented in Figure 1. We showed an important population of CD4^+^ T cells, present in UI skin, which increased as the lesion evolved. Cellularity increased significantly at the CC stage and continued increasing at the PA stage for all cell types except mast cells, whose cellularity peaked at the CC stage (Figure S4). Of note, mast cells accounted for an important population of skin immune cells (13.0 ± 2.8 % of CD45^+^ cells), similar to the proportion of macrophages and resident dendritic cells in UI skin (Figure 1a-c). We observed that some cells such as neutrophils, monocytes and iDCs were scarce at the CC stage and started to be recruited at the PA stage. A massive infiltration of cells classically known to invade tissues early in the inflammatory process (namely neutrophils, monocytes, iDCs and B cells) was also observed at the PA stage (Figure S4).

**Figure 1.**
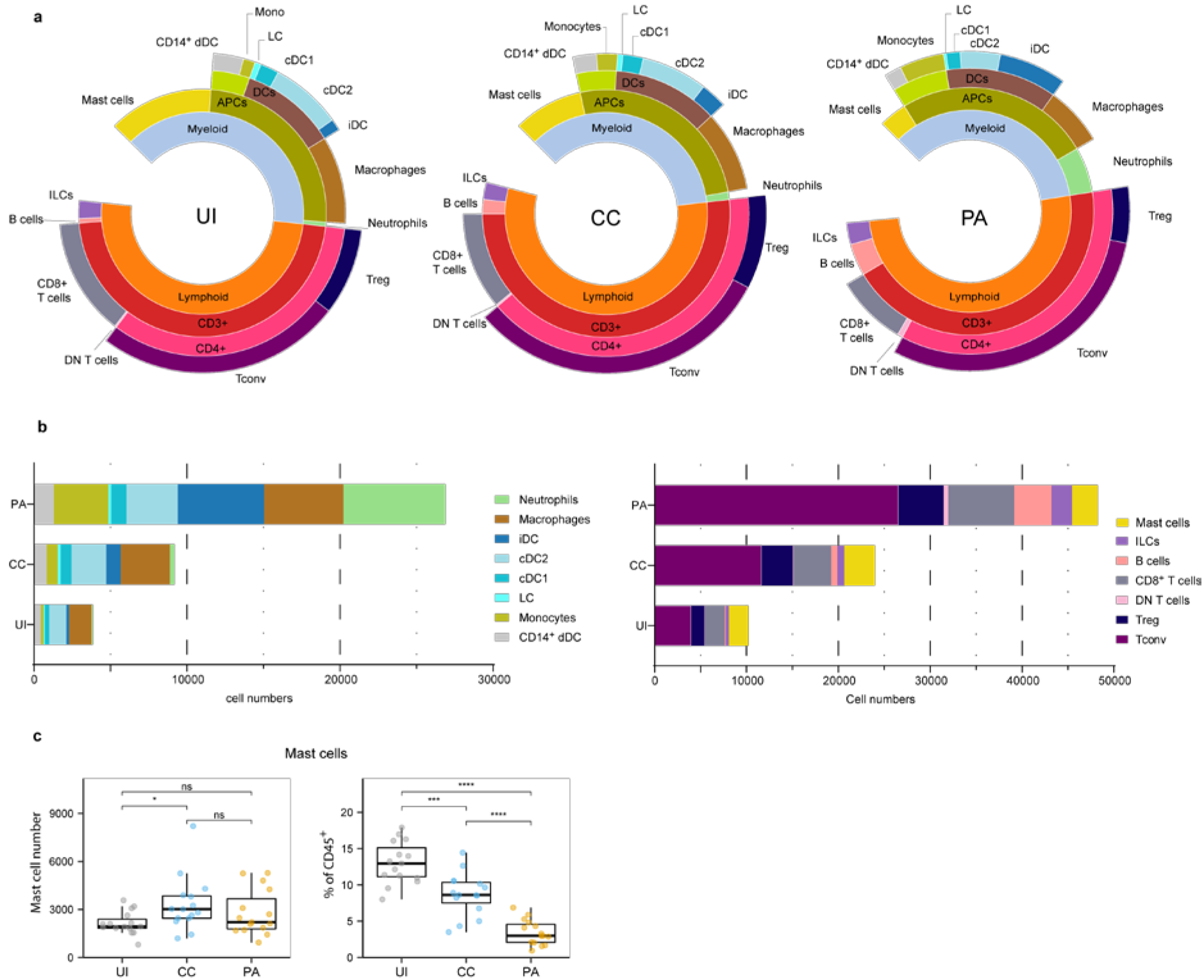
Immune landscape of the early stages of acne. (a) Sunburst representation of the distributions and frequencies of the various cell subsets identified in the 3 stages. Results are expressed as percentages (means) of CD45^+^ cells. (b) Stacked bar graph representing the average number of the major immune cell populations identified with the two flow cytometry panels. (c) Data for mast cell number and percentages of CD45^+^ cells. Friedman test and Dunn’s post hoc tests * p<0.05; ***p<0.001; **** p<0.0001; ns, not significant. The other cell number evolution between the 3 stages are provided figure S4.

We investigated the activation status of the identified professional antigen presenting cells (APCs) by measuring CD40 and CD86 cell surface expression (Figure S5). Because CD86 was expressed by virtually all APCs analyzed, we measured CD86 expression level (CD86 gMFI fold increase over control, Figure S5a-b). We observed that among skin resident APCs, macrophages, cDC2s and CD14^+^ DCs showed high CD86 expression level in UI skin. This expression level remained unchanged in macrophages from CC and PA biopsies while it increased from CC through PA stage in cDC2s. CD40 expression analysis showed a similar pattern (Figure S5c-d). We further analyzed the mast cells that constitute an important sentinel cell population (Figure 1) and observed that the high affinity receptor for IgE, FcεRI, was upregulated through all of the acne stages studied (Figure 2a-b). Moreover, CD69 (an activation marker of mast cells (Gaudenzio et al., 2009, Jayapal et al., 2006)) was strongly expressed in all the analyzed acne stages and its expression peaked at the CC stage, indicating that mast cells are activated very early in the acne process (Figure 2c).

**Figure 2.**
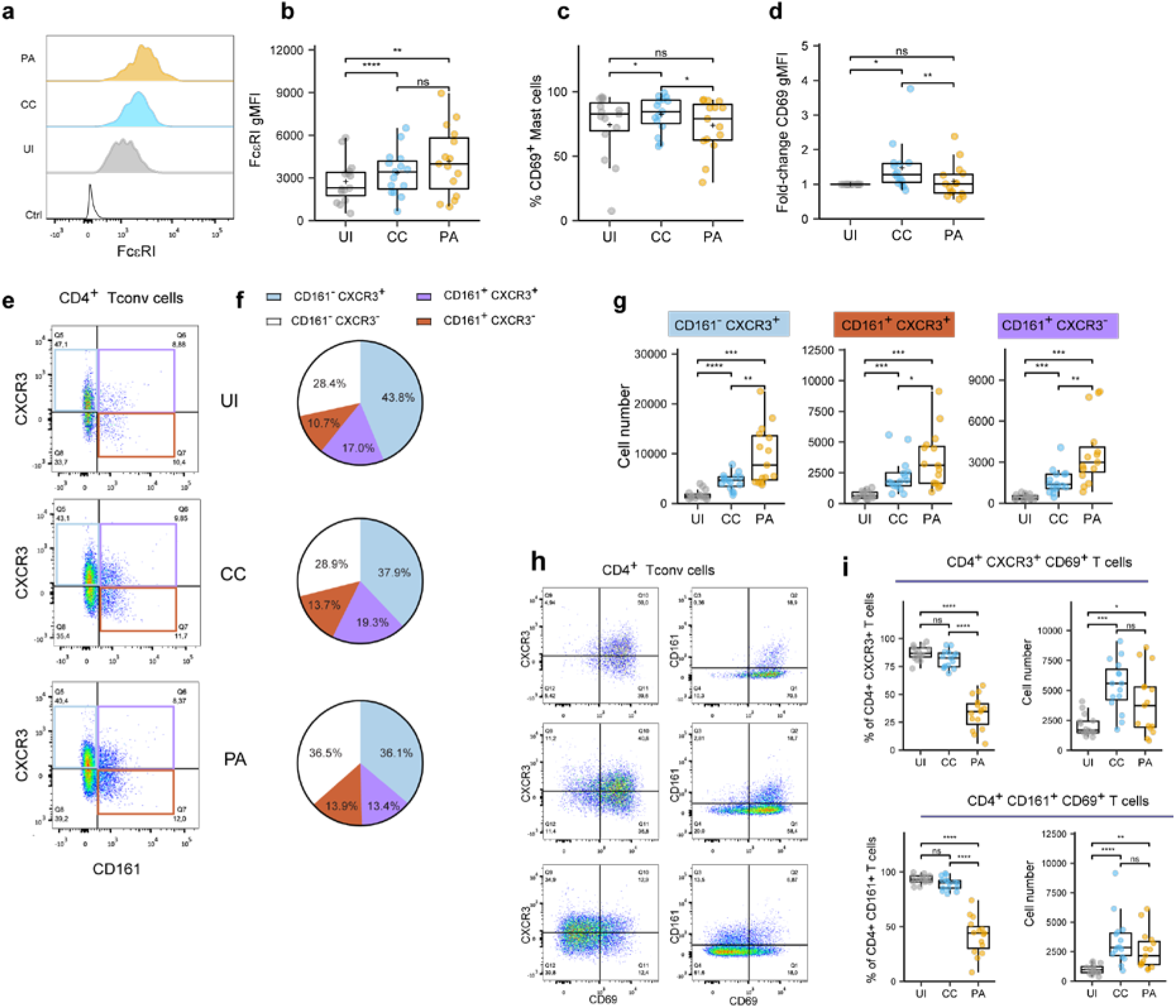
Mast cells and CD4^+^ T cells phenotypes in the early stages of acne. (a-b) FcεRI expression by mast cells. Typical histograms (b) and pooled data as FcεRI gMFI values from the 15 patients. (c-d) CD69 expression by mast cells. % CD69^+^ mast cell (c) and CD69 gMFI fold-change over UI skin (d). Points represent the individual values for the 15 patients. Box and whiskers plots in the style of Tukey. * p<0.05; ** p<0.01; ns, not significant (one-way ANOVA and Fisher’s least significant difference post hoc test). (e) Representative dotplot of CD161 and CXCR3 expression within CD4^+^ Tconv cells (CC stage) and (f) average percentages of CD161/CXCR3 populations within CD4^+^ Tconv cells (15 patients). (g) Percentage of indicated CD161/CXCR3 CD4^+^ Tconv cell populations. (h) Representative dotplot of CD69, CD161 and CXCR3 expression within CD4^+^ Tconv cells (CC stage). (i) Percentage and number of CD69^+^ cells among CXCR3^+^ or CD161^+^ CD4^+^ Tconv cells. Box and whiskers plot in the style of Tukey. Each point represents a patient. Friedman tests were carried out to compare groups and, if significant, were followed by Dunn’s post hoc tests * p<0.05; ** p<0.01; ***p<0.001; **** p<0.0001; ns, not significant.

**Figure 3.**
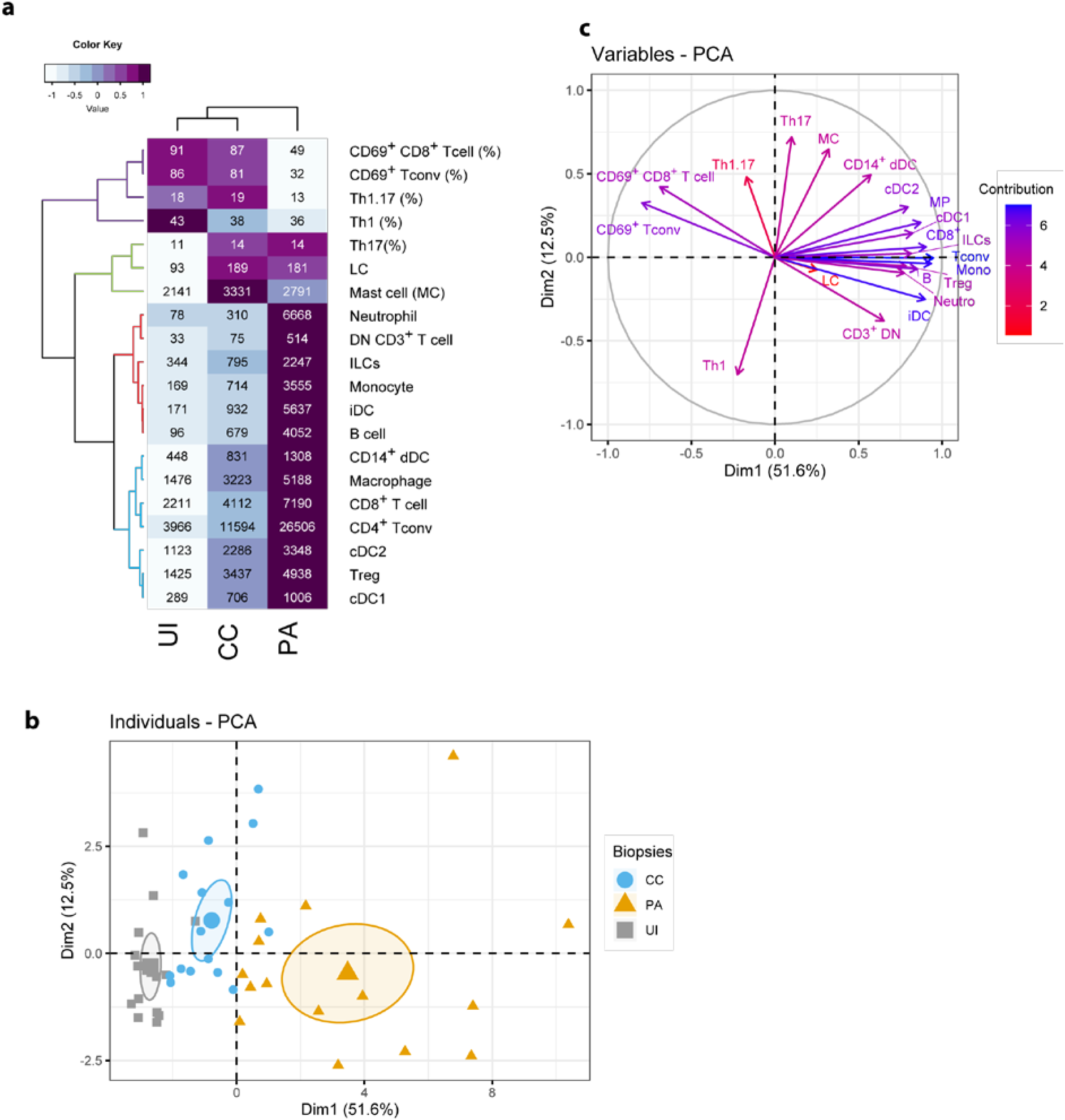
Unsupervised analysis of the skin immune profiles of UI, CC and PA biopsies. (a) Heatmap depicting flow cytometry parameters calculated for UI, CC and PA biopsies (n=15 patients). The mean of each indicated flow cytometry parameter (showed in the cells) was scaled, centered, clustered hierarchically with complete linkage and Euclidean distance measure and visualized in the heatmap using the function heatmap.2 of the gplots R package. Principal component analysis (PCA) was performed with flow cytometry variables shown in the heatmap. (b) Individuals’ PCA. Each dot represents one patient, color was attributed according to the acne stage. The bigger dot shows the group barycenter. The confidence interval of the mean (0.95%) is represented by ellipses. (c) PCA variable correlation plot. Variable contribution is shown by color code. Th1: CD161^−^CXCR3^+^ T conv cells, Th17: CD161^+^ CXCR3^−^T conv cells; Th1.17: CD161^+^ CXCR3^+^ T conv cells; CD3^+^ DN T cells: CD3^+^ CD4^−^CD8^−^ cells.

Because Th17 and Th1 cells have been described as key players in acne, we aimed to identify these subsets by flow cytometry. To cope with the constraints imposed by the limited size of the biopsies, we opted for CD161 and CXCR3 markers to identify Th17 (Maggi et al., 2010) and Th1 populations respectively (CCR6 was not used because it was degraded on the cell surface by the enzymatic digestion step). CD69 was also used as a marker of Ag-experienced skin resident T cells (Mackay et al., 2015, Mueller and Mackay, 2016, Watanabe et al., 2015). CD4^+^ conventional T cell analysis showed that an important proportion of CD161^+^ cells was found in the skin (Figure 2 d-e). It is worth noting that the majority of CD161^+^ Th cells co-expressed CXCR3 in the UI and CC stages. CXCR3^+^ CD161^−^Th cells represented the main Th cell population. All these population exhibited a dramatic increase at the CC stage and kept increasing at the PA stage (Figure 2f). The vast majority of CD4^+^ Tconv cells, either CD161^+^ or CXCR3^+^, expressed CD69 in the UI and CC stages (Figure 2g-h). The percentage of CD69^+^ Tconv cells decreased markedly at the PA stage. CD69^+^ Th cell number peaked at the CC stage. Taken together, these results indicate that CD161^+^ and CXCR3^+^ Th cells dominate in acne skin and that these cells reside in UI skin as CD69^+^ T cells. An important T cell infiltrate, mainly constituted of CD69^−^T cells, is observed as early as the CC stage.

### Unsupervised analysis of the immune cells involved in CC and PA stages of acne

We next carried out an unsupervised analysis of the dataset obtained by flow cytometry. We included in the analysis the absolute cell numbers of the identified populations (Figure 1) and data concerning CD4^+^ Tconv and CD8^+^ T cells (Figure 2) as input variables. The heatmap (Figure 3a) summarizes these data and presents a hierarchical clustering based on their correlation, making conspicuous the grouping of some variables (see colors in the dendrogram). The first group (blue color in Figure 3a) consists of blood leukocytes (neutrophils, monocytes, ILCs and B cells) whose number increased at the CC stage and peaked at the PA stage. These cells are known to be recruited during the inflammatory process or to differentiate after blood precursor recruitment. They participate in the local inflammatory response and contribute to the exacerbation of inflammation. Another group (red color in Figure 3a) consists of skin resident leukocytes, whose absolute number increases substantially at the CC stage (CD14^+^ dDCs, macrophages, cDC2s, cDC1s and T cells). Two other skin resident cells (LCs and mast cells, color green in Figure 3a) increased at the CC stage and stabilized at the PA stage.

Principal component analysis (PCA) showed that the expression profiles of the 3 acne stages are distinguishable (Figure 3b). The first dimension clearly distinguishes the different stages, with the PA stage showing the highest values. Analysis of the loadings plot showed that the number of CD8^+^ T cells, Tconv cells, monocytes, iDCs, and neutrophils (Figure 3c) contributed strongly to this dimension. It indicates that the first dimension is associated with inflammatory cell infiltration. On the contrary, UI biopsies are gathered on the left of this axis following the direction of CD69^+^ T cells. CD69^+^ T cells and low infiltrate characterize the UI skin. Another group of variables contributed to the second PCA dimension (notably Th17, Th1.17, mast cells and CD14^+^ dDCs). The CC biopsies exhibited globally higher values in this second dimension, indicating that these populations are defining features of the CC stage. Moreover, their associated vectors are adjacent in the correlation circle, indicating that these variables are positively correlated (Figure 3c).

### Microscopy analysis of IL-17-producing mast cells in the early stages of acne

Because we identified cDC2 activation during the early stages of acne and because these cells have been shown to promote type 3 responses (Schlitzer et al., 2013), our next approach was to create a cartography of cDC2 and IL-17^+^ CD4^+^ T cells using confocal microscopy. We employed the CD1c marker (Granot et al., 2017, Zaba et al., 2007) to identify cDC2s.

To generate a global view of each biopsy, tile scans of 30-40 sequential z-planes were acquired, creating a 3D mosaic of adjacent image stacks. In the CC and PA stages, we observed a perifollicular accumulation of CD4^+^ T lymphocytes and CD1c^+^ cells (Figure S6a and S7a, areas highlighted in yellow) as well as clusters of perivascular immune cells enriched in CD4^+^ T cells and cDC2s along the rete subpapillare. In some biopsies, clusters of CD4^+^ T cells could be observed deeper in the dermis. CD1c staining revealed a lining of cDC2s just beneath the dermo-epidermis junction in the papillary dermis (Figure S6a and S7a). It was not possible to compare the cell infiltrates between CC and PA stages because cellularity depended upon the area sampled by the slicing procedure. Nevertheless, perifollicular accumulation of CD4^+^ T cells appeared larger in some PA biopsies when compared to their CC counterparts, which was combined with a thickening of the epidermis at the PA stage (Figure S6e). We observed that the majority of CD1c^+^ cells resided either in the CD4^+^ T cell enriched areas or just beneath the dermo-epidermal junction (Figure S6a-b and S7a-b). Unexpectedly, IL-17 staining revealed that CD4^+^ T cells represented only a minor fraction of IL-17^+^ cells, but non-CD4^+^ cells that stained positive for IL-17 appeared to be preferentially located in CD4^+^ T cell clusters. IL-17 staining was often punctate and covered a large part of the cell cytoplasm (Figure S6c-d and S7c-d).

These results prompted us to identify the other cells that produced IL-17. We labelled neutrophils with myeloperoxidase (MPO), macrophages with FXIIIA, mast cells with tryptase and the whole T cell population with CD3 staining. Using triple staining immunofluorescence (tryptase, CD3 and IL-17), we observed an abundance of tryptase/IL-17 double positive cells in CC biopsy sections, indicating that mast cells stained positive for IL-17 (Figure 4a-b). Macrophages did not stain positive for IL-17 (Figure S8). We observed some IL-17^+^ MPO^+^ cells in PA biopsies infiltrated by neutrophils but they were not the main IL-17^+^ cell population (Supplementary Figure S9). We next quantified the CD3^+^/IL-17^+^ cells or tryptase^+^/IL-17^+^ cells in MC and PA biopsies in 4 patients (Figure 4c). Because we have previously shown that mast cells can cooperate with antigen-experienced T cells (Gaudenzio et al., 2009, Gaudenzio et al., 2013), we also analyzed whether the IL-17 positive or negative mast cells were inside or outside the T cell clusters (perifollicular or perivascular). The contingency tables obtained from the quantification of 4 CC and 4 PA biopsies from 4 different patients indicated that mast cells inside the T cell clusters were clearly more prone to produce IL-17 than their counterparts outside T cell clusters (Figure 4d).

**Figure 4.**
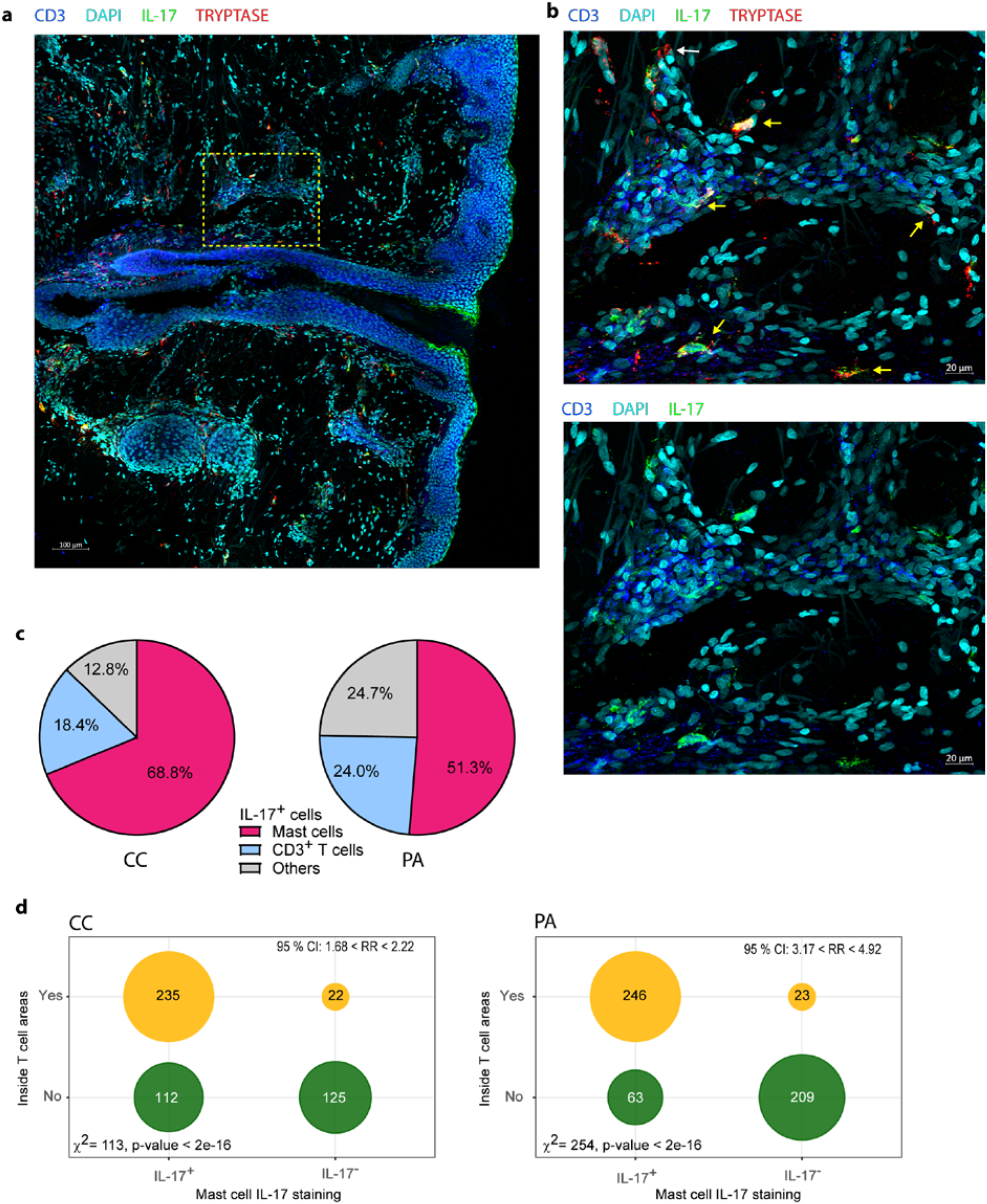
Mast cells are the main cells staining IL-17^+^ in the early stages of acne. (a) A representative image depicting IL-17, CD3 and tryptase staining (CC stage). (b) High-magnification image showing IL-17^+^ mast cells (yellow arrows) near perifollicular area (bottom) and perivascular area (top); white arrow points IL-17^−^mast cell. (c) Image quantification of IL-17^+^ cells among CD3^+^ (T cells), Tryptase^+^ (mast cells), pooled data from 4 patients either at CC or PA stage as indicated. (d) Tryptase^+^ cells were classified according to IL-17 staining and to location (inside or outside T cell rich areas as depicted in Figure S2), resulting contingency tables of counts are plotted (pooled data from 4 patients); χ^2^ test for independence and relative risk are indicated.

### Activated memory CD4^+^ T cells induced IL-17 production by mast cells

We next investigated the stimuli able to induce IL-17 production by mast cells using primary mast cell lines derived from peripheral blood CD34^+^ precursors (hMCs). We first stimulated hMCs with classical activating stimuli (IgE/anti-IgE, substance P, C5a, TLR2 and TLR4 ligands with or without priming with AhR ligands) or combinations of cytokines known to promote IL-17 production in T cells (such as IL-1β, IL-23 and IL-6) but we did not detected any IL-17 production in the conditions tested (data not shown).

The relationship observed between IL-17 production by mast cells and their localization in T cell rich areas prompted us to test whether activated CD4^+^ T cells might induce IL-17 production by hMCs. Moreover, the observation that some IL-17^+^ mast cells located in T cell rich areas were in close proximity to CD4^+^ T cells reinforced this hypothesis (Figure 5a). We cocultured hMCs with *ex vivo* sorted effector/memory CD4^+^ T cells stimulated with anti-CD3/CD28 coated beads. Because both CD4^+^ T cells and mast cells could produce IL-17, we FACS-sorted mast cells after coculture and measured *IL-17A* mRNA by RTqPCR. IL-17A mRNA was induced after 4h coculture and increased up to 24 hours (Figure 5b). Coculture in a transwell system showed that cell-cell contacts are necessary for the activated CD4^+^ T cell to drive IL-17A gene transcription in mast cells (Figure 5c). To analyze IL-17 expression at the protein level, we analyzed mast cell/CD4^+^ T cell cocultures by confocal microscopy following immunofluorescence staining of intracellular IL-17 and tryptase to unambiguously identify mast cells (Figure 5d). Whilst less than 3% of mast cells were IL-17^+^ following coculture with unstimulated CD4^+^ T cells, more than half of them were IL-17^+^ following coculture with anti-CD3/CD28-stimulated CD4^+^ T cells. Collectively, these results indicated that mast cell interaction with activated memory/effector CD4^+^ T cells promoted IL-17 production by mast cells.

**Figure 5.**
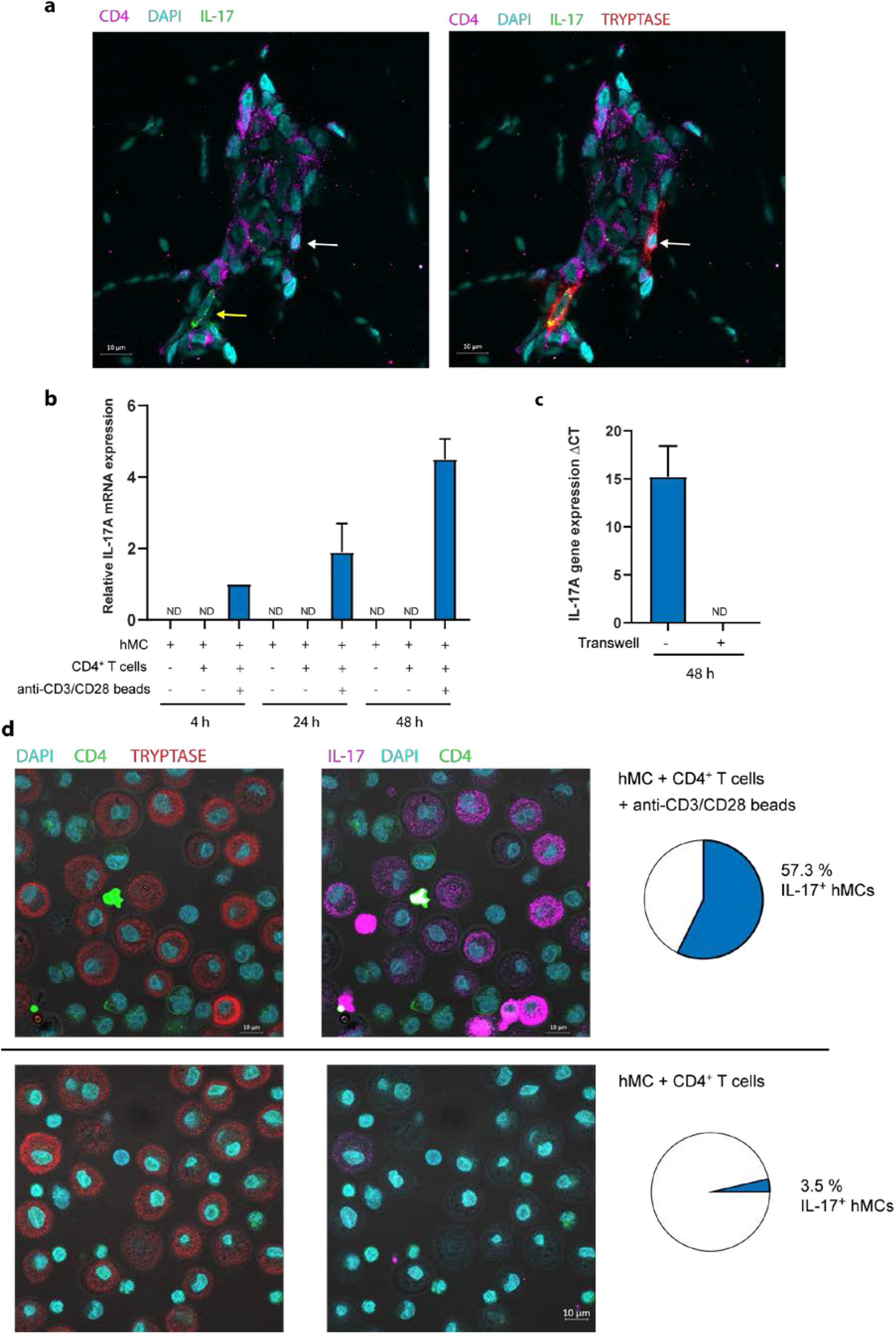
Activated CD4^+^ memory T cells induce IL-17A mRNA expression in mast cells in a contact-dependent manner. (a) Typical example of tryptase+ mast cell in contact with CD4+ T cell in CC biopsy. Yellow arrow shows IL-17^+^ mast cell, White arrow shows IL-17^−^mast cell. (b) hMCs and memory CD4^+^ T cells were cocultured (with or without anti-CD3/CD28 coated beads) for indicated time. hMCs were FACS sorted and IL-17A mRNA expression was quantified by RTqPCR. (c) hMCs and activated memory CD4^+^ T cells with anti-CD3/CD28 coated beads were cocultured for 48h with or without a transwell system. hMCs were FACS sorted and IL-17A mRNA expression was quantified by RTqPCR. Mean +/-SD from 3 independent experiments (hMCs from 3 different donors). (d) IL-17 and tryptase intracellular stainings in hMCs and memory CD4^+^ T cells cocultured (with or without anti-CD3/CD28 coated beads) for 48h. Typical confocal images and quantification of IL-17^+^ hMCs (n>200) representative of 3 independent experiments (3 different hMC/CD4^+^ T cell pairs).

## DISCUSSION

In this study, we employed both flow cytometry- and microscopy-based approaches to delineate the immune landscape of the early stages of acne. Our quantitative analysis corroborated the view that significant cellular changes occur early in acne lesion development, before clinical symptoms appear.

Our unsupervised analysis clearly demonstrated that specific cell populations cluster together and that different clusters dominate at different stages of lesion development. The evolution of these populations is a defining feature of acne immunopathogenesis. Mast cells appeared as pioneer cells in the acne immune response followed by the group of resident APCs (CD14^+^ dDCs, macrophages, cDC2s, cDC1s) and resident T cells. These three groups of cells could be suspected to play a role in the initiation of the acne process. The hypothesis of an immune response burden in uninvolved skin in patients with acne is reinforced by the fact that we observed a high percentage of activated mast cells and macrophages (according to CD86 and CD40 expression) in UI biopsies. This point is reinforced by CD69 expression on the mast cell surface in UI skin as well as during the CC and PA stages, suggesting that these cells are implicated very early in acne. It would have been interesting to analyze UI skin by microscopy but the local ethic committee refused the second biopsy of UI skin.

To analyze the T cell response specifically, we devised a flow cytometry panel to identify CD8^+^ T cells, CD4^+^ Tconv and Treg cells. Because repeated exposure to commensal microorganisms (notably *C. acnes*) is expected to occur in acne and because previous reports showed the importance of the Th1 and Th17 axes in acne (Agak et al., 2014, Kelhala et al., 2014, Kistowska et al., 2015), we chose to target Th1, Th17 and TRM cells in our analysis. We employed CXCR3 to identify Th1 cells (Brodie et al., 2013, Groom et al., 2012, Langenkamp et al., 2003) and CD161 to identify IL-17 producing T cells (Cosmi et al., 2008, Kleinschek et al., 2009). Although CXCR3 and CD161 are often found in Th1 and Th17 cell subsets respectively, they do not strictly define them. Nevertheless, these markers provided a reasonable approximation for our analysis of CD4^+^ Tconv cells. The presence of a substantial contingent of Th17 cells (CD4^+^CD161^+^ Tconv cells) in the skin at steady state is not surprising because these cells are known to participate in immune surveillance of barrier organs. CD4^+^CD161^+^ Tconv cells are expected to belong to the Th17 subset, but CD161 expression is not synonymous with IL-17 production because these cells need to engage their TCR to produce cytokines. Our work quantifies for the first time CD161^+^ T cells in acne and we note a clear increase of Th17 cell number in the CC stage, indicating an early involvement of this subset in acne pathogenesis. In line with previous reports (Agak et al., 2014, Kelhala et al., 2014, Kistowska et al., 2015, Mashiko et al., 2015), we show that Th17 cell numbers are increased in acne and we support the notion that these cells are recruited and/or proliferate during CC formation while they are already present in UI skin of patients with acne. Moreover, the combined analysis of CXCR3 and CD161 showed the presence of cells expressing both markers (about 15% of Tconv cells) that are reminiscent of pathogenic Th17.1 cells (Annunziato et al., 2007, Ramesh et al., 2014), that produce both IL-17 and IFN-γ previously described in autoimmunity or autoinflammation (Stockinger and Omenetti, 2017).

The analysis of CD69 expression by CD4^+^ and CD8^+^ Tconv cells showed that the large majority of these cells were CD69^+^ in UI skin. CD69 is a classical early activation marker *in vitro*. It also appears to be associated with tissue retention *in vivo* thanks to its capacity to counteract S1P-mediated egress and is expressed by memory T cell subsets that are retained in the periphery (Mackay et al., 2015, Mackay et al., 2013). CD69 is expressed notably by tissue-resident memory T cells (TRM) (Kumar et al., 2017, Mackay and Kallies, 2017, Sathaliyawala et al., 2013, Thome et al., 2014, Watanabe et al., 2015) (Klicznik et al., 2019). TRM are now under intensive investigation and CD69 expression appears to be a hallmark of TRM cells. In line with previous reports, we observed that the percentage of CD69^+^ CD4^+^ or CD8^+^ T cells was very high in UI skin biopsies (Klicznik et al., 2019, Wong et al., 2016). It is likely that these cells in UI skin represent TRM cells that accumulated via repeated immune responses against skin commensal microorganisms in patients with acne. The fact that CD69^+^ absolute cell number increased (for both CD4^+^ and CD8^+^ T cells) at the CC stage and plateaued at the PA stage (a non-significant slight decrease was measured) suggests that these TRM cells might be involved and proliferate in the very early steps of acne pathogenesis. At the PA stage, an important recruitment of CD69^−^effector T cells and/or possibly a loss of CD69 by some proliferating TRM cells is observed. This scenario is compatible with what was observed by Park et al, in skin challenged with *C. albicans* (Park et al., 2018). It is tempting to speculate that locally activated commensal-specific TRM cells (in cooperation with local APC such as cDC2s, macrophages or CD14^+^ dDCs) drive the progression of the acne inflammatory process. The early involvement of T cell responses prompted us to investigate by microscopy the location of IL-17 producing CD4^+^ T cells in relation with cDC2s. We focused on cDC2s because they were expected to drive Th17 responses (Collin and Bigley, 2018) and because we observed that they were activated in CC biopsies. In agreement with Natsuaki et al (Collins et al., 2016, Natsuaki et al., 2014), we observed large perivascular clusters of DC and CD4^+^ T cells in CC as well as PA biopsies. Nevertheless, we were surprised to observe that CD4^+^ T cells were not the main IL-17^+^ population and that instead, IL-17 appeared to be produced by large mast cells, notably in CC biopsies. It appears that in acne, as was reported in psoriasis, T cells are not the main population producing IL-17 (Keijsers et al., 2014, Lin et al., 2011).

We found that mast cell number peaked at the CC stage and represented about 10-12% of CD45^+^ cells. This observation might be explained by an increase of stem cell factor (SCF) production by keratinocytes, since alteration of the microbiota was reported to increase the mast cell population indirectly by acting on SCF production by keratinocytes (Wang et al., 2017, Wu et al., 2019). IL-17 is not a typical mast cell cytokine but several studies reported IL-17^+^ mast cells in different pathophysiological conditions (Brembilla et al., 2017, Buckland, 2010, Chen et al., 2016, Hobo et al., 2020, Hueber et al., 2010, Kenna and Brown, 2013, Lin et al., 2011, Liu et al., 2014). However, no mechanism underlying IL-17 production by mast cells in these situations was reported. Here, we showed that activated Th cells license mast cells to activate IL-17A transcription via cell-cell contacts. This indicates that mast cells are able to produce their own IL-17 following cell-cell cooperation with CD4^+^ T cells. This result echoes our previous work showing that Th cell-mast cell cooperation fosters mast cell degranulation upon IgE/Ag stimulation (Gaudenzio et al., 2009).

Taken together, these results show for the first time that mast cells are a major effector of the early immune response in acne, which is controlled by the Th cell response. We can infer that mast cells serve as early IL-17 producers in acne pathogenesis, driven by the nascent Th17 and Th1.17 responses (possibly due to the reactivation of TRM cells). In later stages of the pathology, neutrophils take over and amplify the IL-17 production. This study paves the road for further investigations to evaluate the role of mast cells in acne beyond IL-17 production. Moreover, it identifies mast cells as new strategic therapeutic targets in acne.

## Abbreviations

APC: antigen presenting cell
CC: closed comedone
DC: dendritic cell
hMC: primary human mast cell line
iDC: inflammatory dendritic cell
ILC: innate lymphoid cell
MPO: myeloperoxidase
PA: papule
PCA: Principal component analysis
PSU: pilosebaceous unit
SIS: skin immune system
Tconv: conventional T cells
TRM: resident memory T cell
UI: uninvolved skin

## ACKNOLEDGEMENTS

We thank the flow cytometry and imaging core facilities of the INSERM UMR 1043, CPTP and of the INSERM UMR 1037, CRCT, Toulouse, France.

This research was supported by INSERM (Institut national de la santé et de la recherche médicale) and by Pierre Fabre Dermo-Cosmétique (PFDC).

## AUTHOR CONTRIBUTIONS

Conceptualization: D.R., E.E.; Supervision: E.E., Formal analysis: E.E.; Investigation: Y.E., E.L., L.B., T.N.; Visualization: E.E., E.L.; Validation LG, EE; Writing – Original Draft Preparation: E.E.; Writing - Review and Editing: B.M.

