## Supplementary material for "IL-17^+^ mast cell/T helper cell axis in the early stages of acne": Figure S

### Supplementary Figures

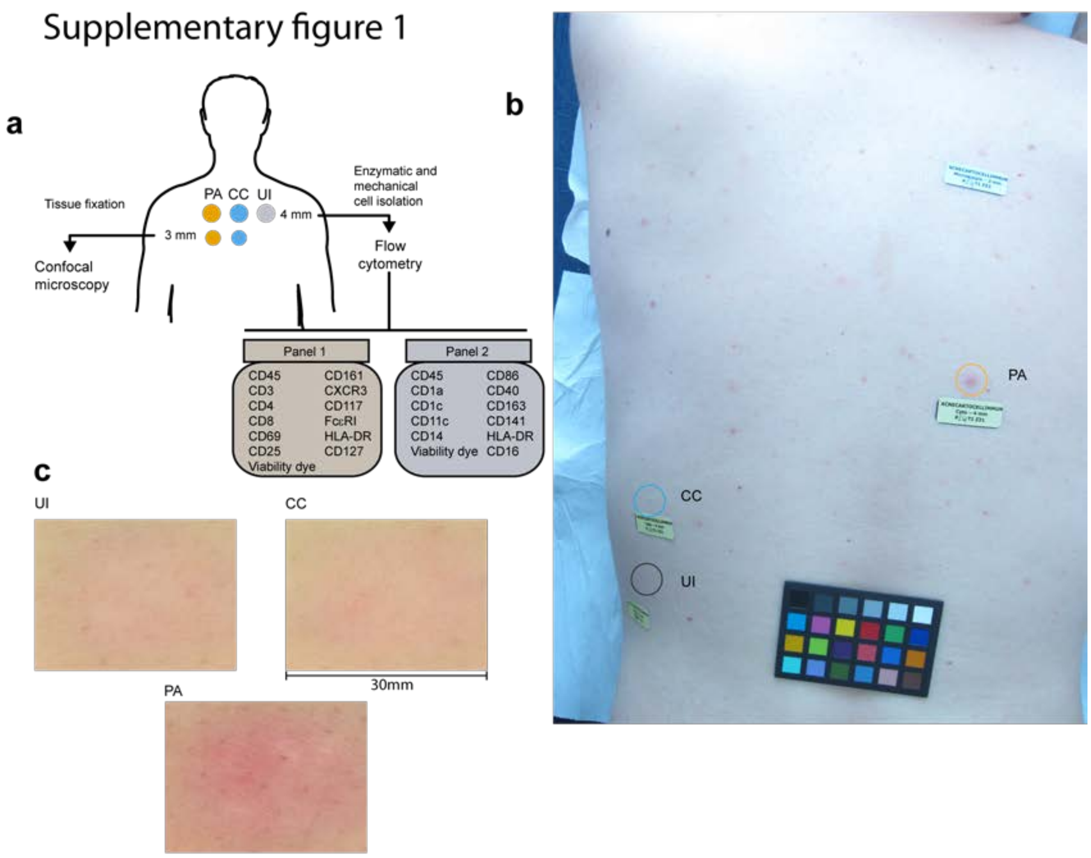


**d**

| Study subject characteristics | |
|---|---|
| Number of subjects | 15 |
| Age, years | 20 to 30 - median = 24 |
| Gender | Male |
| Fitzpatrick skin phenotype | 4 subjects phototype II<br>9 subjects phototype III<br>2 subjects phototype IV |
| Acne severity : IGA score (Investigator's Global assessment) | 3 or 4 |
| Inclusion criteria | At least 2 papula and 2 microcomedones on the back and IGA score >= 3 on the back |
| Facial acne | Mild or absence of facial acne |

**Figure S1. Clinical study characteristics.** (a) Processing workflow for biopsies. The two panels used for immunofluorescence are indicated. (b-c) typical example of areas biopsied for UI skin, CC and PA stages (b) and higher magnification, Pixience dermoscope (c). (d)

Characteristics of the patients involved in this study. CC : closed comedone; PA, papule; UI, uninvolved skin.

### Supplementary figure 2

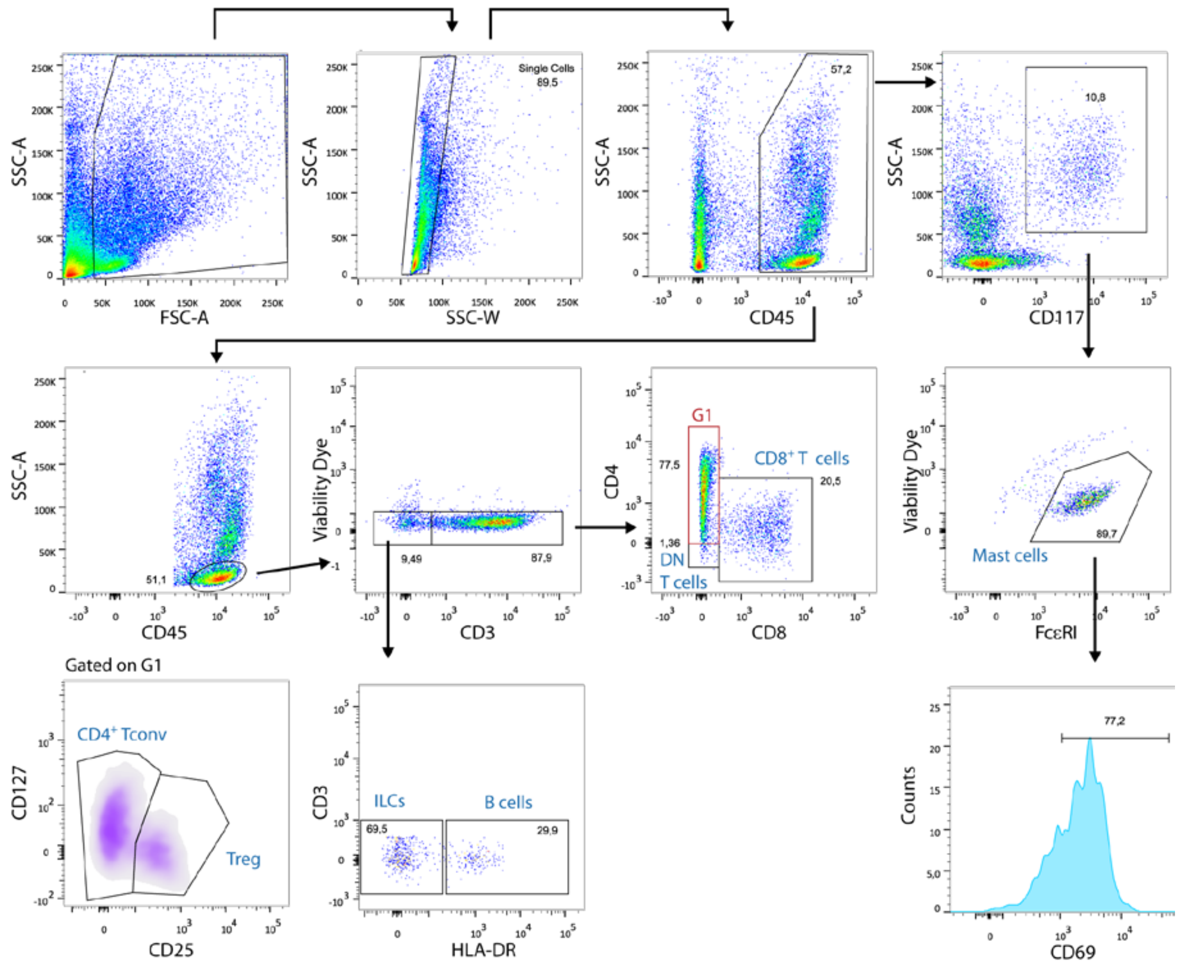


**Figure S2. Gating strategy used for the analysis of skin cells stained with Abs from panel 1.** Representative dotplots and histograms from a CC biopsy. Targeted populations are indicated in blue. Among $CD4^+$ T cells, regulatory T cells (Treg) were separated from conventional T cells (Tconv) as $CD127^{low}$ $CD25^{high}$ cells. Among $CD3^-$ cells, B cells were identified as $HLA\text{-}DR^{high}$ cells and the remaining cells as innate lymphoid cells (ILCs).

Supplementary figure 3

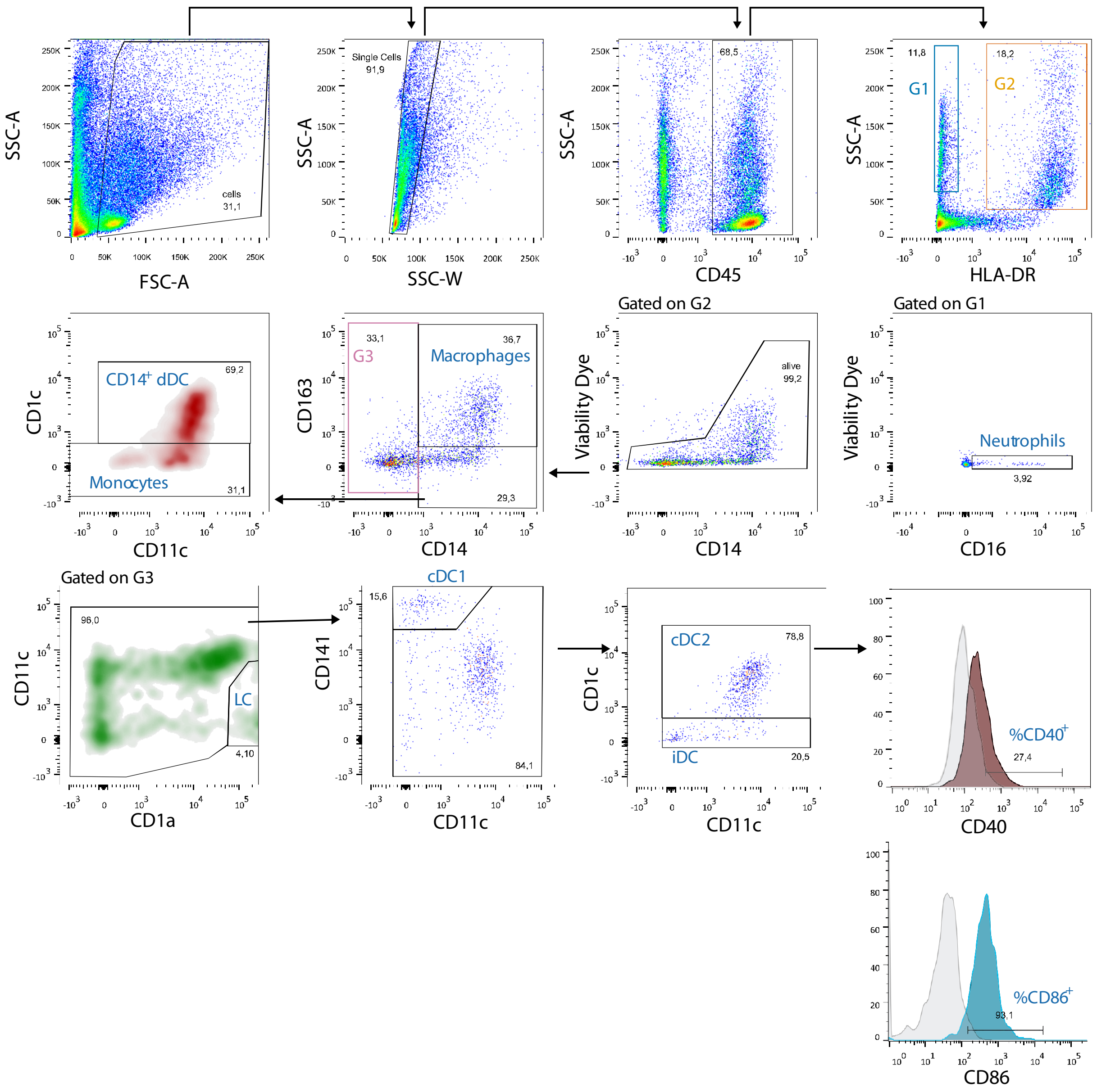


**Figure S3. Gating strategy used for the analysis of skin cells stained with Abs from panel 2.** Representative dotplots and histograms from CC biopsy. Targeted populations are indicated in blue. Among HLA-DR$^{+}$ SCC$^{high}$ cells, macrophages were identified as CD14$^{+}$ CD163$^{+}$ cells (Zaba et al., 2007), conventional dendritic cells as CD14$^{-}$ and among CD14$^{+}$ CD163$^{-}$ cells we used the CD1c marker to separate monocyte (CD1c$^{-}$) from CD14$^{+}$ dermal dendritic cell (CD14$^{+}$ dDC) (Nestle et al., 1993). These latest were shown to be close to monocytes and a transient population of monocyte-derived macrophages (McGovern et al., 2014). Among the CD14$^{-}$ DC population, we identified Langerhans cells (CD1a$^{high}$/CD11c$^{low}$) (Bigley et al., 2015), conventional DC1s (cDC1, CD141$^{high}$ CD11c$^{int}$), conventional DC2s (CD1c$^{+}$, cDC2) and a remaining population of $CD11c^{+}$ inflammatory DCs (iDCs) (Clark et al., 2019, Haniffa et al., 2015, Tang-Huau and Segura, 2019).

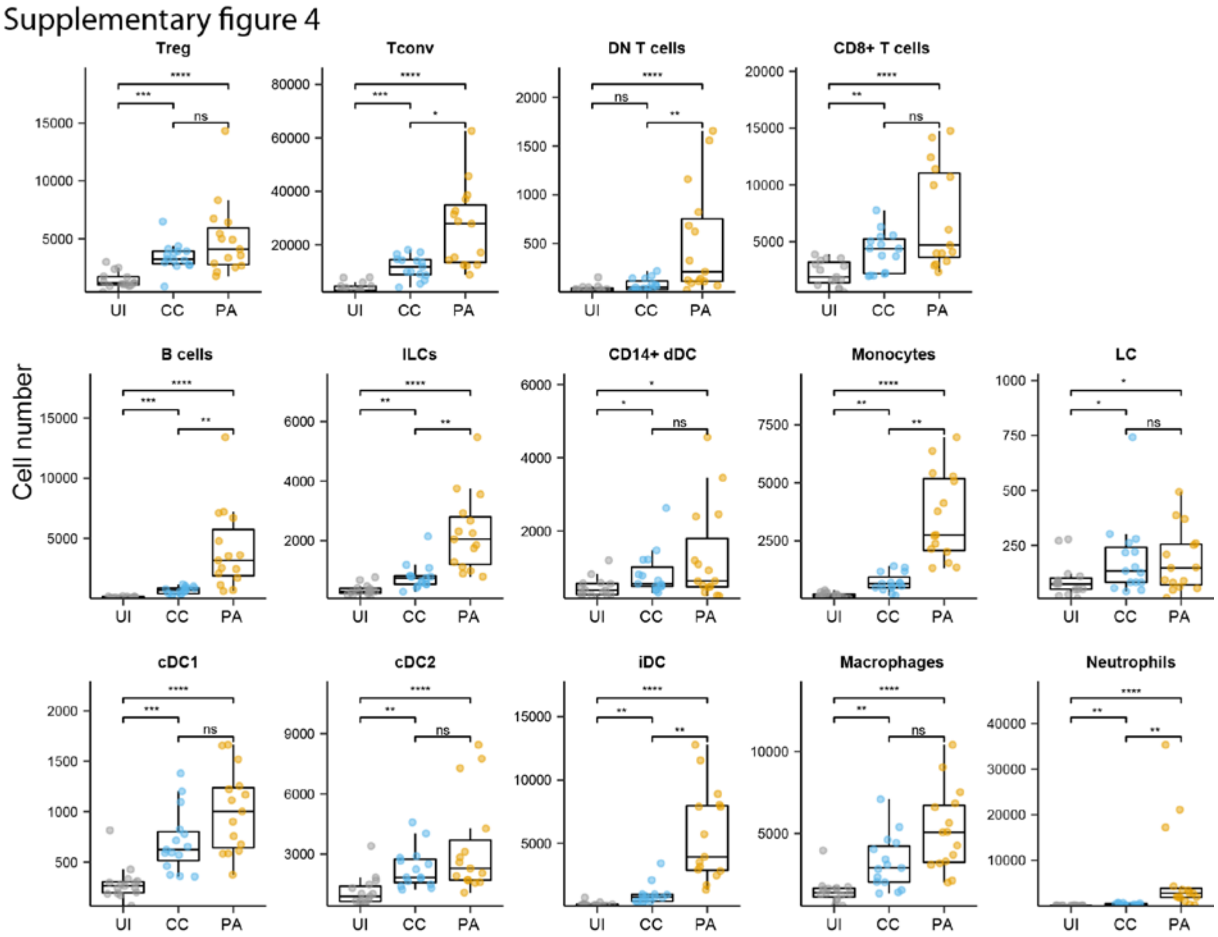


**Figure S4. Absolute numbers of the main leukocyte populations identified by flow cytometry.** Box and whiskers plot in the style of Tukey. Points represent the values for each of the 15 patients individually. Friedman tests were carried out to compare groups and, if significant, were followed by Dunn's post hoc tests * p<0.05; ** p<0.01; ***p<0.001; **** p<0.0001; ns, not significant.

Supplementary Figure 5

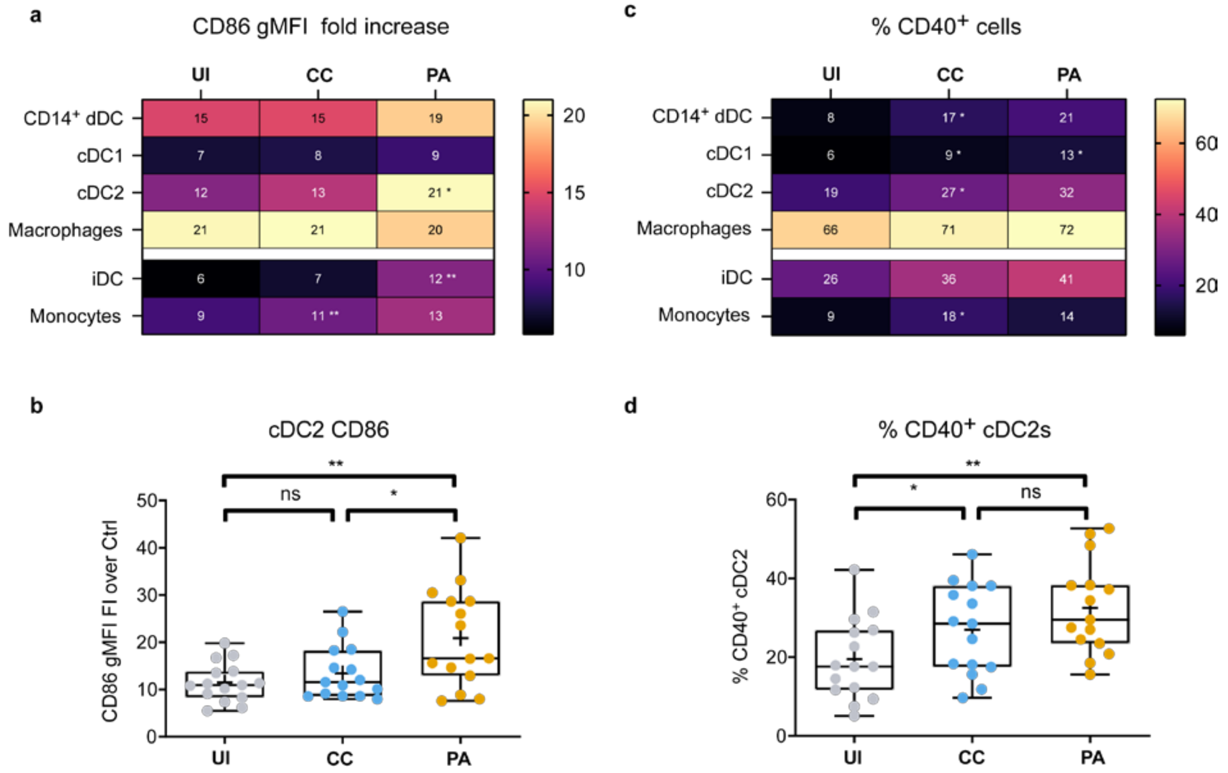


**Figure S5. cDC2 and CD14⁺dDC showed signs of activation during the early stages of acne.** (a) Heatmaps depicting CD86 expression (measured as the fold increase gMFI over the unstained control) (left panel) and the percentage of $CD40^+$ cells (right panel) across each APC population, mean values from the 15 patients are indicated in the cells. Statistically significant differences between UI and CC and between CC and PA conditions are indicated in CC and PA cells respectively, * $p<0.05$, **$p<0.01$. (b) Shown are the cDC2 data from the 15 patients. Box and whiskers represent minimum, $25^{th}$ percentile, median, $75^{th}$ percentile and maximum values of CD86 (c) and CD40 (d) expression by cDC2; points represent the individual values for the 15 patients. The cross (+) represents the mean. * $p<0.05$; ** $p<0.01$; ns, not significant (one-way ANOVA and Fisher's least significant difference post hoc test).

Supplementary figure 6

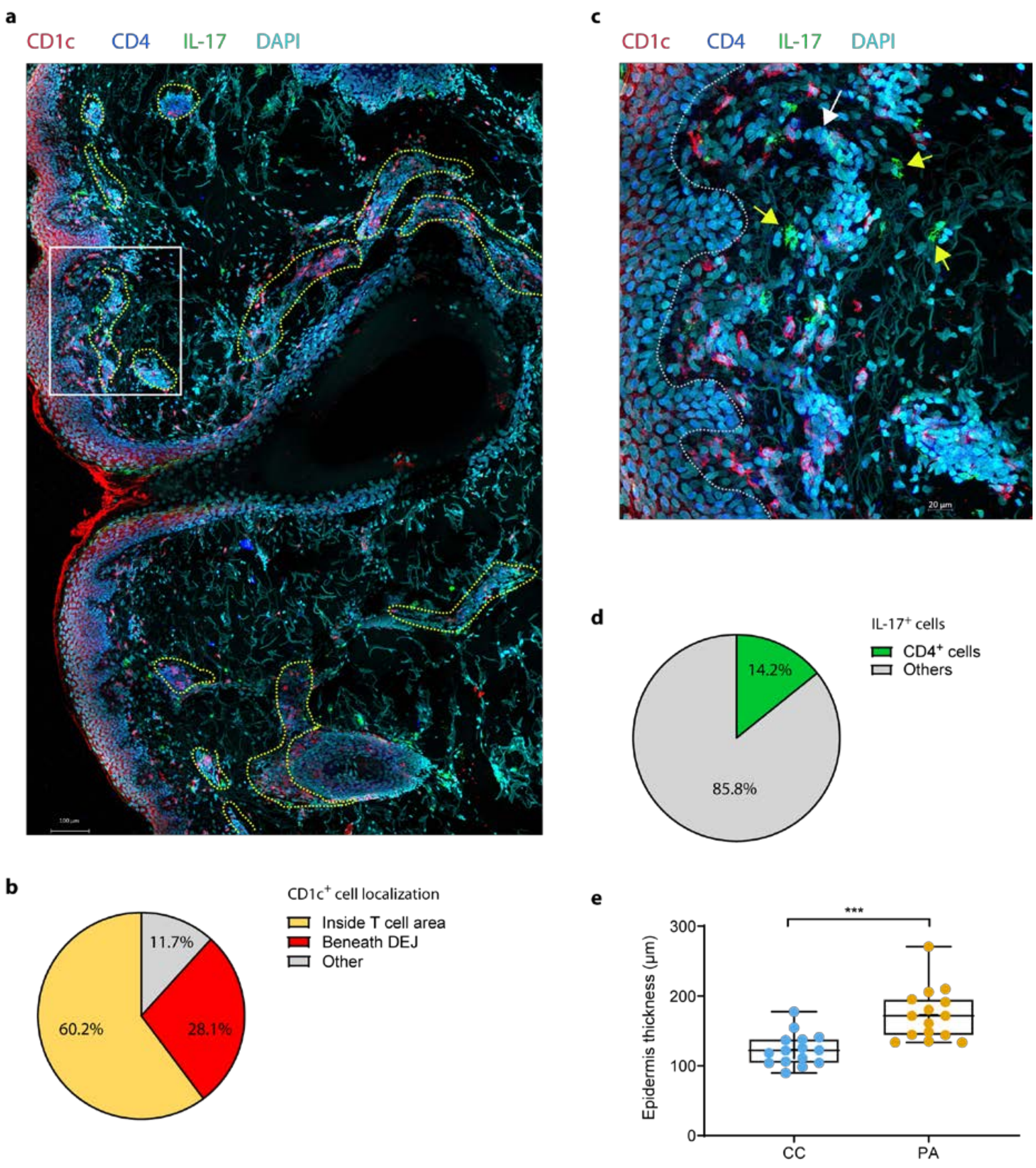


**Figure S6. CD4$^+$ T cells account for only 14% of IL-17-producing cells in CC biopsies.**

(a) Representative confocal laser scanning microscopy tile scan from an entire CC biopsy section presented in maximum intensity projection of a z-stack series. Perifollicular and perivascular regions are delineated with yellow dashed lines. Enrichment of CD1c$^+$ cells beneath the dermal-epidermal junction (DEJ) and inside T cell-rich areas is observed. (b) Quantification of CD1c$^+$ cell localization (n=5 CC biopsies). (c) Magnification of a T cell area showing IL17$^+$ CD4$^+$ T cell (white arrow) and IL-17$^+$ non-CD4$^+$ unidentified cells (yellow arrows). (d) Quantification of IL-17$^{+}$ CD4$^{+}$ cells (3 CC biopsies were analyzed, n=218 IL-17$^{+}$ cells). (e) Epidermis thickness measurement (n=15 patients). Each point represents a patient (mean of 6 measurements taken at regular intervals along the epidermis with Zen software). ***, p<0.001 (Student's *t*-test).

**Supplementary Table S1**

| Antibodies for direct immunofluorescence (flow cytometry) | | | | |
|---|---|---|---|---|
| Antigen | Fluorophore | Clone | Panel | Manufacturer |
| CD3 | BV786 | SK7 | P1 | BD |
| CD4 | PE-cy7 | SK3 | P1 | BD |
| CD8 | PerCP-Cy5.5 | RPA-T8 | P1 | BD |
| CD25 | PE-CF594 | M-A251 | P1 | BD |
| CD69 | BV711 | FN50 | P1 | BD |
| CD117 | APC-R700 | YB5.B8 | P1 | BD |
| CD127 | BB515 | HIL-7R-M21 | P1 | BD |
| CD161 | BV650 | DX12 | P1 | BD |
| CXCR3 | APC | 1C6/CXCR3 | P1 | BD |
| HLA-DR | PE | G46-6 | P1 & P2 | BD |
| FceRI | e450 | AER-37 | P1 | eBiosciences |
| CD1a | APC | HI149 | P2 | BD |
| CD1c | BB515 | F10/21A3 | P2 | BD |
| CD14 | PE-CF594 | MfP9 | P2 | BD |
| CD40 | BV421 | 5C3 | P2 | BD |
| CD45 | APC-H7 | 2D1 | P1 & P2 | BD |
| CD86 | BV711 | 2331 (FUN-1) | P2 | BD |
| CD163 | BV650 | GHI/61 | P2 | BD |
| CD11c | BV785 | 3.9 | P2 | Biolegend |
| CD16 | AF700 | 3G8 | P2 | Biolegend |
| CD141 | PE-Cy™7 | M80 | P2 | Biolegend |
| Primary antibodies for indirect immunofluorescence (microscopy) | | | | |
| Agntigen | Isotype | Clone | | Manufacturer |
| Tryptase | Mouse IgG1 | AA1 | | Millipore |
| IL-17 | Goal polyclonal | | | R&D systems |
| CD1c | Mouse IgG1 | 5B8 | | Abcam |
| CD4 | Rabbit | | | Sigma-Aldrich |
| CD3 | Rabbit | | | Dako |

| | | | | |
|---|---|---|---|---|
| MPO | Rabbit | | | Dako |
| FXIIIA | Rabbit | | | Thermo |

Table S1. Antibodies used in immunofluorescence.

Supplementary figure 7

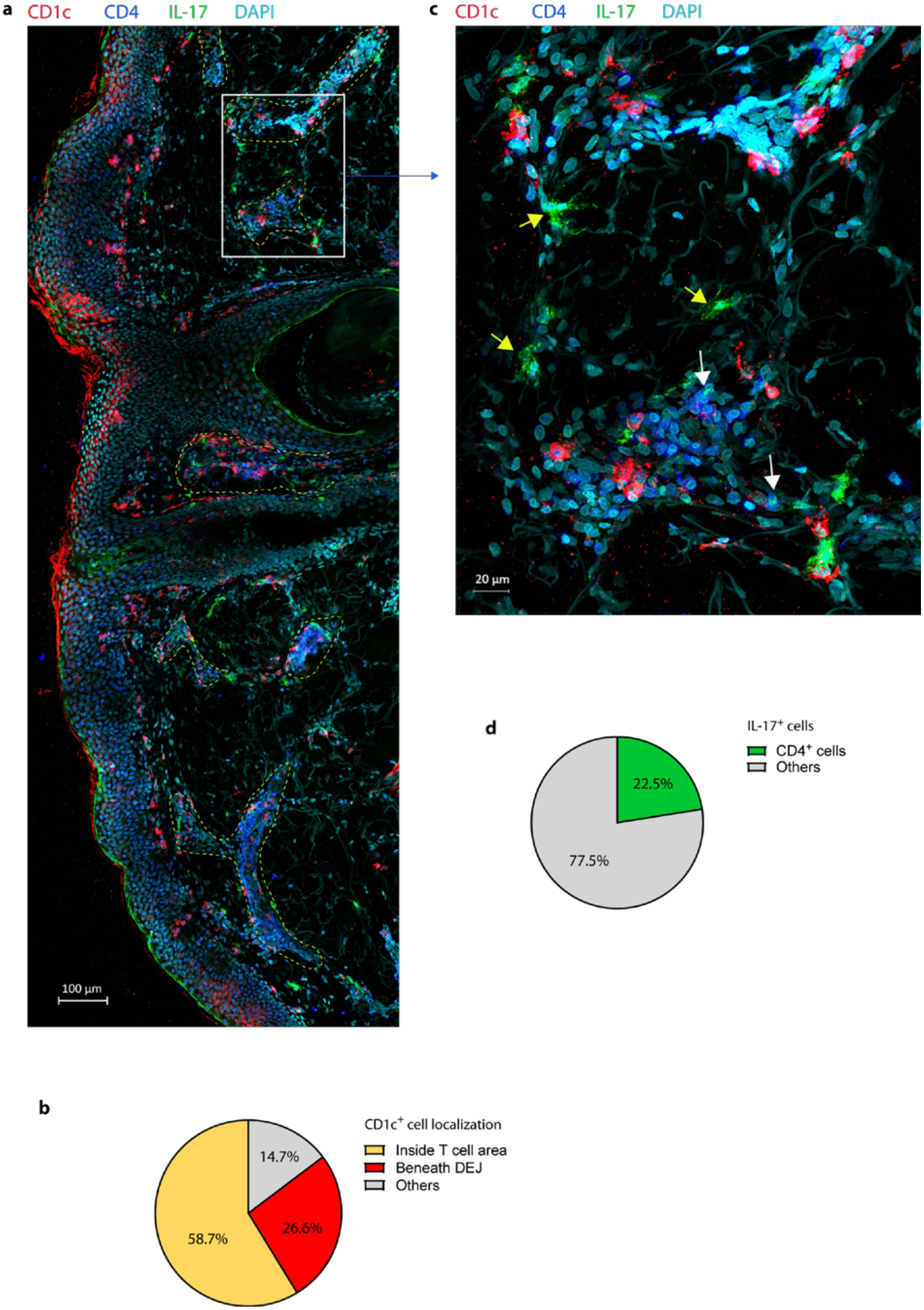


**Figure S7. CD4$^+$ T cells account for less than 23% of IL-17-producing cells in PA biopsies.**

(a) Representative confocal laser scanning microscopy tile scan from an entire PA biopsy section, presented in maximum intensity projection of a z-stack series. Perifollicular and perivascular regions are delineated with yellow dashed lines. (b) Quantification of CD1c$^{+}$ cell localization (n=5 PA biopsies). (c) Magnification of a T cell area showing IL-17$^{+}$ CD4$^{+}$ T cell (white arrows) and IL-17$^{+}$ non-CD4$^{+}$ unidentified cells (yellow arrows). (d) Quantification of IL-17$^{+}$ CD4$^{+}$ cells (3 PA biopsies were analyzed, n=298 IL-17$^{+}$ cells).

### Supplementary figure 8

Tryptase FXIIIA DAPI IL-17

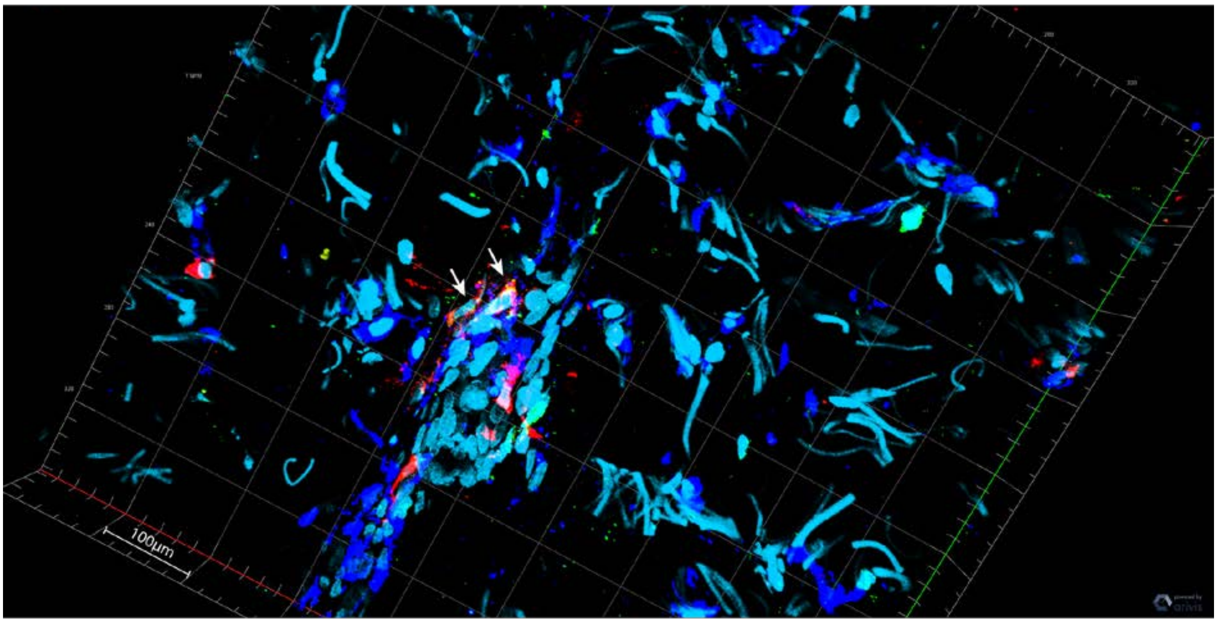


Tryptase FXIIIA DAPI

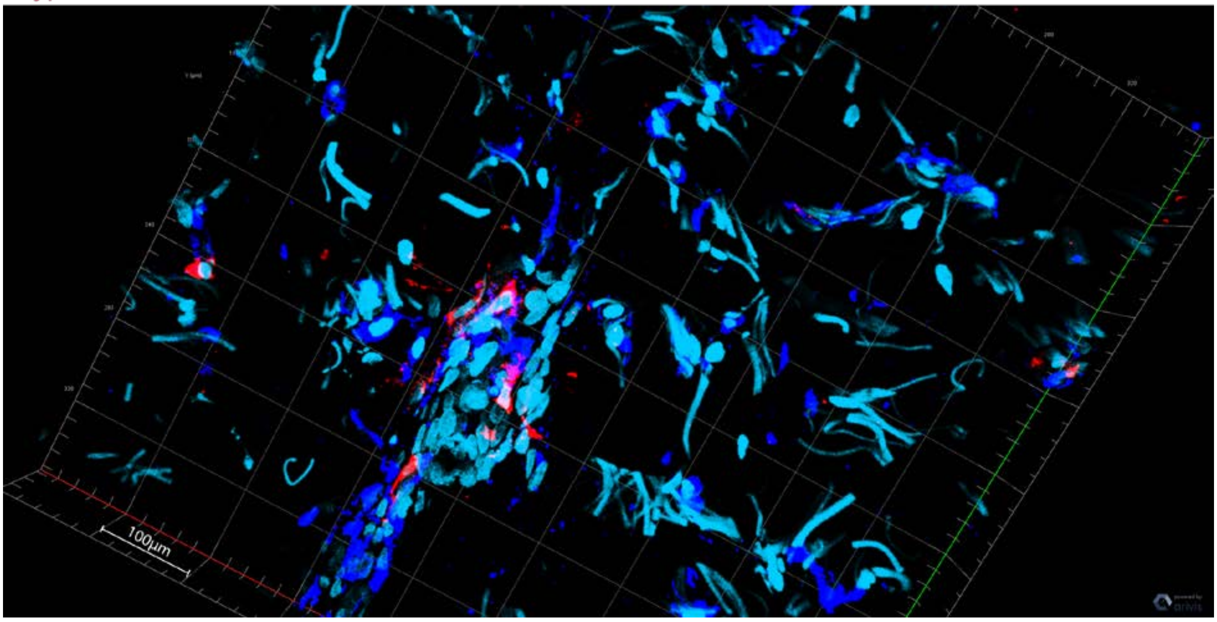

**Figure S8. Macrophages did not produce IL-17 in acne biopsies analyzed.** 3D reconstruction from laser confocal scanning z-stack depicting a perivascular area in the dermis of a PA biopsy. Representative image showing several FXIIIA$^{+}$ (blue) macrophages staining negative for IL-17 (green).

Supplementary figure 9

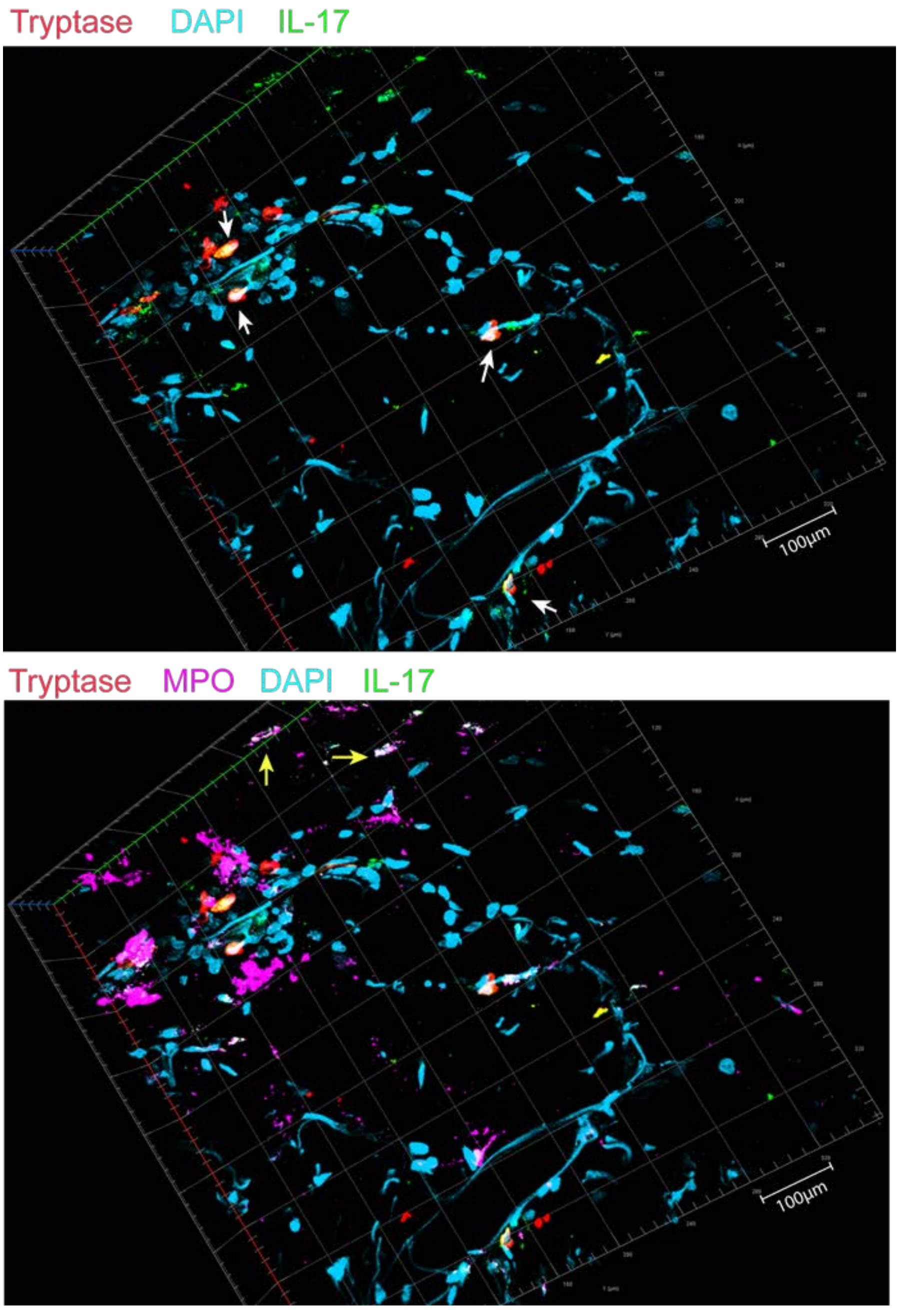


**Figure S9. A fraction of the neutrophils stained IL-17$^{+}$ in PA biopsies.** 3D reconstruction from laser confocal scanning z-stack depicting a perivascular area in the dermis of a PA biopsy.

Representative image showing some $MPO^+$ (pink) neutrophils (yellow arrows) staining positive for IL-17 (green) and some IL-$17^+$ $tryptase^+$ mast cells (white arrows).
